# The expendable male hypothesis

**DOI:** 10.1101/473942

**Authors:** Siobhán M. Mattison, Robert J. Quinlan, Darragh Hare

## Abstract

Matriliny is a system of kinship in which descent and inheritance are conferred along the female line. The theoretically influential concept of the matrilineal puzzle posits that matriliny poses special problems for understanding men’s roles in matrilineal societies. Ethnographic work describes the puzzle as the tension experienced by men between the desire to exert control over their natal kin (i.e., the lineage to which they belong) and over their affinal kin (i.e., their spouses and their biological children). Evolutionary work frames the paradox as one resulting from a man investing in his nieces and nephews at the expense of his own biological offspring. In both cases, the rationale for the puzzle rests on two fundamental assumptions: (i) that men are always in positions of authority over women and over resources; and (ii) that men are interested in the outcomes of parenting. In this paper, we posit a novel hypothesis that suggests that certain ecological conditions render men expendable within local kinship configurations, nullifying the above assumptions. This arises when (i) women, without significant assistance from men, are capable of meeting the subsistence needs of their families; and (ii) men have little to gain from parental investment in children. We conclude that the expendable male hypothesis may explain the evolution of matriliny in numerous cases, and by noting that female-centered approaches that call into doubt assumptions inherent to male-centered models of kinship are justified in evolutionary perspective.

---

> “…for some females there exist important advantages for male care … For many females male parental care has small or negligible effects on female reproductive success, suggesting that as a general explanation for social monogamy, the Male Care is Essential Hypothesis is inadequate.” – Gowaty 1996 [1]

## 1.0 Introduction

Matriliny exists throughout the animal world, but is often considered problematic in humans. The problem created by human matriliny is exclusive to males bound by norms that require them to invest more in their sororal nieces and nephews than in their own children. Such avuncular investment violates a fundamental expectation from Hamilton’s rule [2] that, all else equal, altruism (investment in another individual at some short-term cost to oneself) should be directed toward more closely related kin. Several solutions have been proposed to what has been dubbed the ‘matrilineal puzzle’ [3], based on i) paternity certainty, ii) the differential impacts of resources on male versus female reproductive success (RS), and iii) other considerations affecting the benefits of avuncular support of sororal nieces and nephews. In this paper, we present the expendable male hypothesis, which questions fundamental ethnographic and evolutionary premises of the matrilineal puzzle. We propose that well-known behavioral principles defined by theories of parental investment and cooperative breeding can explain differential male investments in their own and in others’ children. Our fundamental claim – which must be empirically validated – is that it is seldom adaptive for males to invest in nieces and nephews *to the detriment* of their own biological children. Rather, ecological, economic, and social factors reduce the benefit of paternal effort for mothers and their children [4], generate poor returns to male investment in pair-bonds and paternal investment, biasing behaviors toward female control of resources and more peripheral roles for males.

We present our argument as follows. First, we provide a definition of matriliny that emphasizes cooperation among females. Next, we review relevant animal literature on female philopatry and male allo-care, in which we find no analogs to support the existence of a matrilineal puzzle in broad comparative perspective. We examine some cross-cultural trends in matriliny including its expressions within cultural systems that prescribe patrilineal organization (i.e., “matrifocality”). We offer critical examination of dominant hypotheses for matriliny including the roles of paternity certainty and daughter-biased investment. These earlier models offer promising leads that we use to elaborate the evolutionary ecology of matriliny in the expendable male hypothesis, in which matriliny results from environments that favor female control of resources and limited male investment in offspring.

### 1.1 Defining matriliny

Matriliny is ambiguously defined [5,6], variously incorporating notions of genealogical descent, corporate descent (i.e., descent operating to join related individuals into a collective body (e.g., a lineage) exercising rights and governing institutions), inheritance, post-marital residence (“locality”), and female-biased cooperative kinship networks. In non-human animals (hereafter, “animals”), these domains of kinship mostly overlap (e.g., matrilineal inheritance is found alongside female philopatry), but this is not always the case in humans (see Surowiec and Creanza, this issue). We thus advance a definition that may be applied across contexts and taxa. Specifically, we define matriliny as a *system of behaviors that bias investment toward matrilineally related kin* [see also 7]. Our definition includes inheritance of resources, rank, title, or information and other forms of cooperation that are biased toward matrilineally related kin. The focus on observable behavior [8–10] separates our definition from more ambiguous metrics of matrilineality such as corporate descent, which arguably apply only to humans and, while affording greater opportunities for certain forms of cooperation among relatives, do not preclude alternatives [e.g., 11]. Finally, our definition requires observation of *actual* behaviors as opposed to stated norms. Kinship behaviors frequently contradict stated norms [12,13] and natural selection acts on behaviors – not stated norms – because enacting norms through behaviors is what generates differential fitness outcomes.

## 2.0 Matriliny in Comparative Perspective

### 2.1 Explanations of Matriliny in Animals

> “Female philopatry enhances the potential for cooperative relationships to arise through kin selection, and may thus facilitate the development of social bonds and the formation of social groups.” – Silk 2007:540 [14]

Understanding the expression and correlates of matriliny in animal societies can shed light on the specific features of human matriliny that emerge as difficult to explain. Matriliny is the predominant form of social organization among mammals. However, its expressions are highly variable to the point that no single explanation for its existence emerges as most likely. We were unable to find published evidence of significant avuncular investment in nieces and nephews (let alone to the detriment of a male’s own offspring), which suggests that it is avuncular investment – not matriliny per se – that is most difficult to explain in human social organization.

Matriliny as defined above is common among gregarious animals [15]. For example, ongoing affiliative relationships or territorial overlap among female kin arise in cetaceans, canids, ungulates, primates, birds, and rodents. Female philopatry, where females tend to remain in their natal communities while males disperse, occurs across taxa and is predominant among mammals: Trochet et al. [16] recorded male dispersal in 77 of the 111 (69%) mammal species they reviewed. Such arrangements lead to aggregations of group-living or neighboring females, and create opportunities for related females to direct benefits towards each other. Moreover, matrilineal behavior takes many forms across species. Matrilineal killer whales *(Orcinus orca)* and African elephants *(Loxodonta africana)* transmit social and ecological information with high fidelity through natalocal matrilines, including significant inputs from post-reproductive females [17,18]. An adult cheetah *(Acinonyx jubatus)* female ceases to interact with her mother while continuing to reside on and eventually inheriting her mother’s territory [19]. Diverse matrilineal species, from farm cats *(Felis catus)* [20], to lions *(Panthera leo),* spotted hyenas *(Crocuta crocuta)* [21] and African elephants [22], engage in communal suckling, whereby lactating females feed related and unrelated young. These different expressions of matriliny suggest that the costs and benefits of matrilineal sociality are variable and warrant empirical and theoretical attention [see also 23].

Yet the causes of matriliny, per se, have been little explored in ethology. Rather, causes must be inferred from arguments concerning the factors selecting for female philopatry, on the one hand, and the benefits of sociality, on the other. For example, an influential model of primate sociality [24] posits that female philopatry and nepotistic relationships allow co-resident females to better defend patchily distributed resources, whereas female dispersal is favored when resources are more evenly distributed. Variants of this ‘socioecological model’ have been extended to other mammals [25] and to incorporate additional ecological factors selecting for or against sex-biased philopatry and sociality. Such factors include better detection and evasion of predators; increasing genetic diversity to reduce risks of inbreeding; effects on pathogen exposure and infection; and reducing risks of infanticide from unfamiliar males [26–28]. In all cases, strong biases in favor of female philopatry and nepotistic relationships among females are attributed fundamentally to the inclusive fitness benefits (i.e., personal fitness benefits and fitness benefits to relatives) derived from living with kin [14]. Why such benefits are discussed in relation to *female* sociality isn’t made explicit in these arguments, but many are likely derived from the expectation that social and dispersal decisions among female mammals are predicated fundamentally on the distribution of resources whereas male decisions center on the distribution of females [25,29].

By defaulting to a female perspective, such causal arguments potentially conflate female philopatry with nepotistic relationships among philopatric females. In particular, it is not clear in these explanations whether the proposed causal factors – resource distribution, inbreeding avoidance, etc. – favor female philopatry, nepotistic female cooperation, or nepotistic female cooperation *given* female philopatry. Moreover, costs and benefits of philopatry and dispersal extend to both males and females, so it is unclear why these explanations focus exclusively on females. We suspect that this may be motivated by the widespread occurrence of female philopatry among mammals [15,29,30], which may render matriliny relatively ‘uninteresting’ to explain. A bipartite theory that builds matriliny from first principles – i.e., that considers the factors favoring female (or natal) philopatry as well as female sociality – would provide clarity, even if such an endeavor may not be simple. There remains little consensus as to which factors (e.g., predation, inbreeding) favor male-versus female-biased dispersal [16,31]. Moreover, female philopatry and nepotistic relationships may be mutually reinforcing, creating infinite regress when considering how these features coevolve to produce an integrated socioecological system of matriliny.

Regardless, female-biased philopatry is neither necessary nor sufficient to cause matrilineal social organization. Numerous animals are female-philopatric and not necessarily matrilineal, for example mountain lions *(Puma concolor* [32]) and wolverines *(Gulo gulo* [33]) in which solitary females hold territories but do not actively cooperate with each other frequently. Moreover, matrilineal social organization is seen in species with different dispersal patterns [16]. Natal philopatry (aka natalocality, in humans) also provides opportunities for female kin to continue affiliative associations [5,30,34] and matriliny has been reported in natal philopatric species, including canids, felids, viverrids, hyanids, and cetaceans [15]. Even female-biased dispersal does not prevent matrifocal kinship behaviors. For example, although male-biased dispersal is more common in non-human primates, related females often emigrate together to nearby populations or otherwise build female-centered affiliative alliances within new groups [35,36]. Moreover, dispersing females might provide benefits to their female relatives and those relatives’ offspring by reducing competition over space, mates, or food [37]. Finally, rates of dispersal vary considerably within systems of sex-biased dispersal [25,31], affording the opportunity for building and maintaining matrilineal alliances, even in species where such alliances are not normative [35]. Thus, matrilineal kinship, if facilitated by female philopatry, arises across dispersal systems and, as the default mode of kinship among mammals, apparently is not particularly problematic^1^.

Because the heart of the controversy in human studies of matriliny concerns the relative costs and benefits of avuncular versus paternal support of juveniles (see 2.2, “Explanations of Matriliny in Humans”), we note here limited scholarship investigating avuncular investment in nieces and nephews in animals. The absence of such scholarship would be easy to chalk up to dispersal patterns if a single sex reliably dispersed prior to maturity, but both natal philopatry [30] and delayed dispersal [40] open the scope for avuncular investment in nieces and nephews in animals. Indeed, in a number of well described matrilineal species, adult males provide allocare [15,16]. Yet, while common, male allocare seems to be restricted to care of siblings or to their own putative offspring [39,41]. Even the male-female “friendships” that have been associated with non-paternal care of offspring in some primates may provide benefits to males through accruing status or eventual access to mates [e.g., 42–44]. In short, whereas avuncular support of nieces and nephews is widely reported in human societies, we were not able to find clear direct comparisons in mammalian societies, matrilineal or otherwise.

We also note a striking difference in the extent to which research on humans versus animals casts matriliny as problematic versus neutral or beneficial. In particular, much ethnographic and evolutionary scholarship views human matriliny as problematic^2^, particularly regarding men’s investment in sisters’ children rather than in their own. By contrast, much of the animal literature treats matriliny neutrally or describes benefits made possible by stable matrilineal structures. Such benefits include female longevity and accumulated social and ecological knowledge [18,46,47], cumulative culture [e.g., 17], infant survivorship [48], and even decreased aggression [30, see also 5]. Matriliny is not sufficient on its own to generate such benefits, yet the stability provided by female-centered kinship may be more likely to generate the conditions to reap the benefits of longevity than more unstable structures associated with looser kinship generated by bonds among patri-kin [49]. The multi-generational structure apparent in some mammalian matrilineal species may also facilitate cultural complexity.

Indeed, eusociality is present only in matrilineal societies (e.g., eusocial insects and naked mole-rats *(Heterocephalus glaber));* levels of cooperation are so high in some of these species that they have been dubbed “superorganisms” to reflect the exceptional degree to which individuals’ interests overlap [50].

Diverse expressions of matriliny throughout the animal kingdom suggest that human matriliny could result from female cooperation to exploit specific types of resources or to protect themselves from various environmental hazards (e.g., predators, infanticidal males). Our review shows further that cooperation among females is not constrained by presumed ancestral male philopatry [35,36]. To the contrary, we can easily imagine a scenario in which related females of a chimpanzee/bonobo-like ancestor dispersed together to some novel territory, eventually joined by an unrelated male or males with relatively limited interest in controlling day-to-day female behavior [see also 36,51]. In chimpanzees [ibid.] and foragers [e.g., 52–54], individuals can and do change their place of residence throughout their lifetimes. Many married Himba women visit their natal group to take advantage of relatives’ support during critical periods of child rearing, suggesting that normative virilocality does not fully constrain female kin coresidence [52]. Adult female chimpanzees sometimes return to their natal group if they fail to establish themselves in a new group following attempted dispersal [55,51]. And even though bonobos *(Pan paniscus)* are male philopatric [56], unrelated females form coalitions to punish aggression and maintain social cohesion within groups [57]. Such behavioral flexibility casts doubt on simplistic generalizations about residence patterns -- for example that forager groups are always patrilocal, female chimpanzees always leave their natal group, or males always exercise control in patrilocal species -- and point instead to context-dependent behavior. More generally, the pervasiveness of female-female cooperation in mammals [15,48] supports our contention that matriliny under a wide range of conditions evident from a broad taxonomic perspective is not particularly puzzling.

### 2.2 Explanations of Matriliny in Humans

> “…matriliny is well adapted to any situation in which competing demands for men are higher than demands for material resources.” – Douglas, 1969, p.130

Female-biased kinship is uncommon but recurrent in human societies: the Standard Cross-Cultural Sample reports matriliny as the institutional mode of descent in 17% of societies and uxorilocal residence (residence with or near the wife’s kin) constitutes 16% of the societies surveyed in the Worldwide Ethnographic Atlas [58]. The extent of overlap between uxorilocal residence and matrilineal descent is considerable: 70% of matrilineal societies are also uxorilocal (see Surowiec and Creanza, this issue). These statistics belie the complex expression of matrilineal behavior, however. For example, matrilineal inheritance does not proceed via a single route of transmission: resources may be inherited directly from mother (or parents) to daughters, as among the matrilineal Chewa of Africa [59] or, as among 8-9% of cases in the Standard Cross-Cultural Sample, from mother’s brother to sister’s son [7]. Cooperation among kin related through females is also much more extensive than the above would suggest. Viri-(residence with or near the husband’s kin) and neo-(residence in newly established domicile) locality do not preclude investment in children by matrilateral kin in WEIRD (Western, Educated, Industrialized, Rich, and Democratic) [60,61] or small-scale societies [62]. Indeed, many societies default to matrifocal social organization even under stated norms of patriliny [e.g., 63–66].

It is therefore surprising that while matriliny is considered evolutionarily neutral or beneficial (maybe even unremarkable) among ethologists, it is frequently considered problematic in studies of humans [e.g., 3,67–70]. Although some ethnographers have emphasized the problems solved by matriliny [e.g., 71, see also 69] “it has [more frequently] been customary to draw attention to the difficulties of working a matrilineal system”, which, among kinship systems is a “cumbersome dinosaur” [69]. Dubbed the ‘matrilineal puzzle’ by Audrey Richards [3], a particular problem surrounds the allocation of male authority, which, according to Richards, is officially granted to men over their natal households (i.e., because they fill the important role of mother’s brother), but simultaneously exerted by a man over his spouse and children [72–74]. This problem does not affect male authority in patrilineal kinship systems, because men’s authority over their natal kin, spouse, and children overlap as in-marrying women are incorporated into their husband’s patrilineage (an in-marrying man is socially inferior to his wife’s brother and not ordinarily considered part of his wife’s lineage following marriage in matrilineal systems [3]). *Prima facie,* matriliny is also puzzling to evolutionary scholars where it involves disproportionate avuncular investment into matrilateral nieces and nephews^3^ [7,34,45,59,67,75–77]. Because nieces and nephews are genetically only half as related to a man as are the man’s biological children, such a pattern of investment defies the logic of Hamilton’s rule, which predicts, all else being equal, greater investment in more closely related kin [2].

Evolutionary attempts to ‘solve’ the puzzle have invoked three main considerations: i) the effects of paternity certainty on the allocation of male parental investment; ii) sex-based differences in the rate at which resources are translated into reproductive success; and iii) other considerations affecting the benefits of avuncular support of sororal nieces and nephews. We describe each of the proposed solutions in turn, and outline a fourth solution – the expendable male hypothesis – which suggests that men in matrilineal societies invest relatively little in any children, often direct their energy elsewhere (e.g., toward mating) and that men’s effort on behalf of children and households is highly substitutable; i.e., kinswomen can compensate for any deficits in men’s contributions. Table 1 summarizes these hypotheses and several associated predictions.

Paternity uncertainty represents the oldest and most common explanation of matrilineal inheritance and cooperative behavior. Alexander [78] was the first evolutionary anthropologist to describe what is, in fact, ‘one of the oldest hypotheses in social science’ [67] in evolutionary terms. The adage, “Mommy’s baby; Daddy’s, maybe” captures the source of a fundamental difference between male and female reproduction in mammals: for mammalian females, genetic parentage is virtually assured, whereas there is nearly always room to doubt a male’s parentage. The paternity certainty hypothesis posits that, if a male’s parentage is very insecure, he may do better in terms of inclusive fitness to invest in his sororal nieces and nephews, to whom genetic relatedness is relatively secure [7,76,78–82]. The level of uncertainty necessary to produce the conditions under which men are more related to their nieces and nephews (i.e., the ‘paternity threshold’ [76,81]) ranges from a probability of paternity (P) of 0.268 [81] to 0.33 [79,82] in the short-term, to 0.46 if the compounding geometric effects of paternity on relatedness over several generations are considered [67], to 0.5 if assumptions about the nature of cuckoldry are relaxed [76]. Such levels are almost certainly unrealistically low [12,45,59,76], all falling well below published estimates of paternity in human societies [75,83,84]. The paternity uncertainty hypothesis, if correct, would require its own explanation accounting for the socioecological factors leading to multiple mating in females [85]. Further, more recent theoretical work has cast doubt on whether the paternity threshold model is biologically realistic, and by relaxing model assumptions show that matrilineal inheritance could evolve under any level of paternity certainty [7,76]. The expendable male hypothesis, which we present below, offers a different solution: low paternity certainty is acceptable in matrilineal systems where women are relatively free to make mate choice decisions based on criteria other than male provisioning [see also 86]. Low paternity certainty in matriliny, hence, may be an artifact of expendable males rather than a cause of limited paternal investment in children.

A second class of explanations incorporates the differential impact of subsistence on male versus female RS. Following the logic of the Trivers-Willard hypothesis [87], the matriliny-as-daughter-biased-investment (MDBI) hypothesis [59] posits that parents should direct wealth toward the sex that is most capable of translating wealth into reproductive success. Because female RS has a lower ceiling compared to male RS and because RS is typically more variable in males than in females [88, but see 89], the slope of the RS-wealth relationship is often steeper for males (Figure 1A [90]). Although each unit of investment yields higher marginal inclusive fitness returns via male offspring under this scenario [91], parents who are of low socioeconomic status and whose sons are likely to fail to reproduce gain absolutely more from investing in daughters [e.g., 90, see also 92]. Many forms of wealth are thought to result in steeper fitness gains through males [93] – both livestock and productive land are usually more beneficial to males than females, for example because males are able to use such resources to attract additional mates [45,59,94]. However, some resources may not be particularly conducive to steep gains in reproductive success in either sex; in which case, the hypothesis predicts, paternity uncertainty in sons’ offspring renders it beneficial to invest in daughters (Figure 1b).

**Figure 1.**
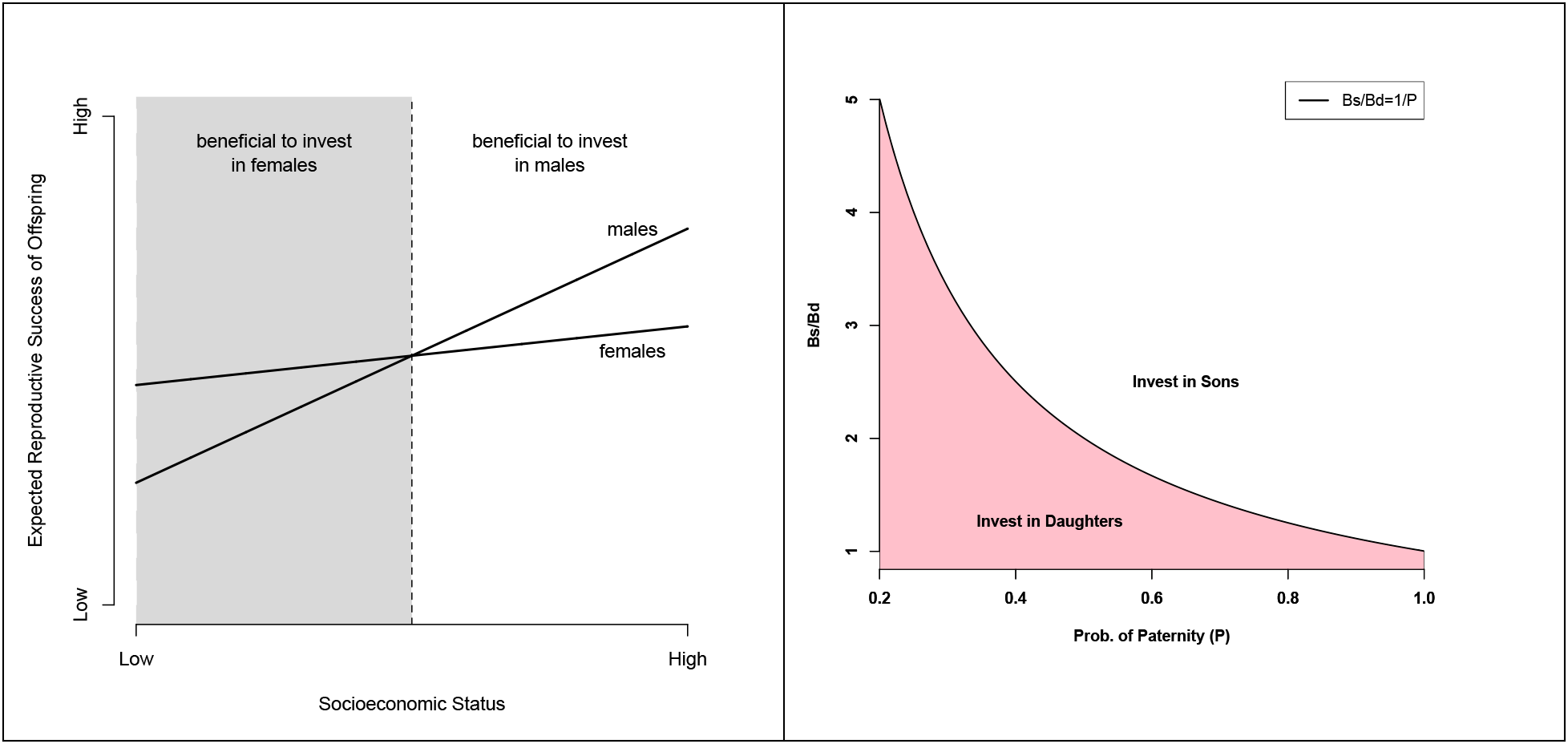
Investments in daughters versus sons. A. Trivers-Willard. The fitness ceiling is lower for females than for males. According to the Trivers-Willard hypothesis, under certain circumstances, this leads to steeper marginal gains to fitness from resources to males. In such cases, poorly resourced parents may benefit from investing in daughters whereas richly resourced parents may benefit from investing in sons (redrawn from [90]). **B. Matriliny as daughter-biased investment (MDBI)**. The MDBI hypothesis incorporates the differential effects of resources on male and female fitness shown in A and the effect of paternity certainty to predict when parents will invest in daughters (matriliny) versus sons (patriliny). The area in pink shows that parents will invest in daughters whenever the additional benefit of investing in sons (Bs/Bd) does not compensate for risk of non-paternity in sons’ offspring (redrawn from [45] based on [59]).

Although two studies of matriliny have described results that support the MDBI hypothesis [45,59] and numerous other findings are consistent with subsistence being crucial to the evolution of sex-biased kinship, by focusing on vertical transmission between parents and their children, the MDBI hypothesis in fact fails to solve the most vexing problem in the evolution of matrilineal kinship – i.e., avuncular investment in sororal nieces and nephews [45]. More recent hypotheses have modeled the ‘father versus uncle’ problem directly, incorporating paternity certainty, the probability that two siblings share the same father [76], the effects of subsistence on inheritance, and additional considerations such as the effects of polygyny and polyandry [7] and the number of communally breeding sisters in a man’s natal household [34], to describe additional benefits that males might accrue via investments directed toward natal kin. Theoretical solutions to the puzzle that do not invoke paternity certainty focus on how the functional form (diminishing versus linear) linking yield to labor affects the returns associated with investing in nieces and nephews over one’s own children [7,95].

We offer a fourth ‘solution’ (section 4.0) that proposes that, at least in some instances, the matrilineal puzzle as implied by ethnographic reports of normative behaviors does not, in fact, exist, and that men’s investments in children follow straightforward evolutionary predictions based on parental investment and cooperative breeding theories.

## 3.0 Problematic Assumptions of the Matrilineal Puzzle

The matrilineal puzzle relies on two assumptions that, if nullified, invalidate the basis of the puzzle: first, that men are in positions of authority over women; and second, that men are interested in the outcomes of parenting. For behavioral ecologists, the problem surrounds avuncular investment in sororal nieces and nephews – transmission of resources (e.g., inheritance) from mother’s brother to sister’s son contradicts the key expectation that a man should invest in more closely related kin. We tackle each of these issues in turn.

### 3.1 Male Authority

> “There is a further difficulty that in most societies authority over a household. is usually in the hands of men.” – Richards, 1950: 246.

Ethnographic material on matrilineal societies suggests that ties of affiliation through females (e.g., ‘suckled at the same breast’) and relatively limited involvement of fathers is frequently normative [e.g., 3,69,96,97]. Although one might expect this to render men’s roles somewhat peripheral, extant literature suggests instead that men wield authority not as fathers, but as mother’s brothers. Among the matrilineal Bantu societies, for example, “.authority passes to the dead man’s brothers or to his sisters’ sons, or to the sons of his maternal nieces.” [3] and “.in most societies authority over households, or a group of households, is usually in the hands of men, not women, as are most of the important political offices” [3]. Flinn (1981) documented a number of compatible examples, including the mother’s brother acting as head of household among the Djuka (citing Kahn 1931), wealth and social position being inherited from maternal uncle to nephew among the Trobrianders (citing Malinowski 1961), and the mother’s brother holding the greatest obligation to provide support and assistance to children among the Plateau Tonga (citing Colson 1961).

Many social scientists have questioned the premise of universal male authority and have emphasized the authority wielded by women in the domestic sphere [e.g., 98–103]. Moreover, even in extended family avunculate households, stated norms stipulate that only one brother of the sibling set act as household head, leaving most men in subordinate positions within households. Furthermore, as behavioral ecologists, it behooves us to ask whether a normative emphasis on male authority plays out in actual patterns of behavior and, perhaps more crucially, what *variation* in behavior tells us about how the *distribution* of ‘authority’ reflects sex biases in cooperation. We contend that, at least in some cases, male authority is nominal rather than real and that the ethnographic and normative emphases on male authority overlook a more stable and central importance of women to provisioning and childrearing [see also 101]. For example, Mattison [45] argued that the nominal transmission of resources from mother’s brother to sister’s son among the matrilineal Mosuo obscures *de facto* transmission down the female line; men’s authority over such resources amounts to usufruct access to land and household resources during their lifetime. The Mosuo also reveal significant heterogeneity in who claims to be household head (户主): male household heads accounted for only ~25 per cent of households residing in areas adhering to a more ‘traditional’ matrilineal lifeway [11]. Moreover, the phrasing of questions in her study had a significant impact on whether male or female authority was emphasized; when Mattison asked who *generally* was the most important decision-maker in a household, the answer was nearly always the mother’s brother; when she asked who was most important in a person’s *own* household, answers varied, but females were commonly listed as heads of house. This underscores that even very careful ethnographic reports of stated norms may reveal inconsistencies, or misrepresent actual patterns of behavior.

For comparison, in Caribbean communities, norms or cultural models invariably emphasize male-headed households and patrilineal inheritance, when actual household organization is often strongly female-centered[64–66]. Women (even married women) exert substantial control over household resources, and inheritance is highly variable [104–108]. Men can and do head households and control family resources in some Caribbean families; however, greater gender-specific risks for Caribbean men can make many men unattractive as mates with potentially poor returns on women’s investment in conjugal relationships [55; see also Macfarlan et al., this issue]. Hence, by our definition, Caribbean matrifocality may be a form of latent matriliny that suggests the likelihood of the *prevalence* of matrilineal behavior in numerous societies nominally adhering to conflicting kinship norms [see also 109]. The Caribbean case further highlights the difficulty of using cultural “traits” (such as patrilineal descent) as the basis of cross-cultural comparison in the study matriliny [e.g., 110]. The behavioral ecological focus on actual behavior provides a means of understanding the evolutionary costs and benefits of female-centered kinship at higher resolution than does reliance on “ideal culture,” often the focus of classic ethnography and cross-cultural work. Although norms and institutions may be products of selection and constrain individual decisionmaking [111], including kinship behavior [67,76], we focus on actual patterns of investment as a more tractable and surprisingly under-studied component of the matrilineal puzzle.

### 3.2 Male Investment

Even in the absence of resource control, individuals can invest time and energy in other individuals. Violation of the first assumption is thus not sufficient to nullify the matrilineal puzzle, so we must also address the extent to which men invest in sororal nieces and nephews versus their own biological children. Whereas the majority of evolutionary work on matriliny takes for granted that men invest significantly in their sororal nieces and nephews, we propose that many men in matrilineal and matrifocal societies may have little interest in childrearing and provide correspondingly low investments therein. If so, this would call into question the second premise of the matrilineal puzzle. Relative to most mammals, male provisioning of offspring in humans is extensive [e.g., 112,113], if also remarkably variable [4]. Male provisioning is correspondingly central to a number of influential theories explaining the evolution of human adaptive complex, which posit that large brains, lengthy juvenile dependence, and nuclear families were made possible and resulted from hunted meat being provided by males to sexually exclusive female partners and their children [summarized in 114]. The observation that men frequently invest in children should obviously not imply that they always do so [e.g., 115,116], yet it seems that there has been an uncritical acceptance of this premise in defining the matrilineal puzzle in evolutionary terms. A quote from Hartung [67] illustrates the problem. In setting up the premise for the matrilineal puzzle, he writes: “Relative to most male mammals, human males give their offspring an exceptional amount of parental investment (Trivers 1972). Accordingly, husbands risk investing a significant component of reproductive success in other men’s children if wives engage in extramarital sex.” Alexander’s [79] solution to the matrilineal puzzle also focuses on misallocated male investment: “.lowered confidence of paternity causes a man’s sister’s offspring to assume increased importance **as recipients of nepotistic benefits**.” (emphasis added). Direct male parental care is generally rare among mammals: Trochet et al.’s review [16] found no instances of exclusive male parental care, and some degree of male participation in only 33 species (30%).

Yet, what if sometimes males in matrilineal kinship systems act like drones in honey bee colonies [117]? That is to say, what if the ethnography does not line up with actual behavior [12] and men are, in fact, doing relatively little to contribute directly to the welfare of their biological offspring or to that of their sororal nieces and nephews? The association between matriliny and protracted male absences (e.g., due to fishing [118], warfare [e.g., 119], or historical involvement in major trade routes [120,121]) means that men are frequently not in a position to invest directly into their households, natal or affinal. Mattison’s ethnographic work among the matrilineal Mosuo concurs with prior descriptions of this population as one with a significant imbalance in female and male workload. In particular, although men engage in occasional heavy labor and obligations related to ritual and political office, women do the vast majority of day-to-day work, including planting, harvesting, and childcare, and also serve as village heads and in other governmental and non-governmental positions [122,123]. Indeed, the limited systematic evidence that we are aware of that addresses workload using behavioral observations is consistent with a lopsided sexual division of labor: Wu et al. [34] report lower levels of observed farming behavior by Mosuo males than by Mosuo females and by Han (majority, patrilineal Chinese) males. He et al. report that “Mosuo males are allowed to reside with their mothers and sisters, who feed them while they do relatively low levels of household domestic or agricultural labor” [124]. Similarly, unmarried men in matrifocal Dominica spend about 17% of their daylight time working (manual labor and childcare) compared with 25% for married men, 50% for women, and 26% for girls 11 to 15 years old [125]. This evidence points to the need to verify empirically the contributions that men make to their natal and affinal households in specific cultural contexts – if male authority and investment in the natal household is limited, then the basis for the matrilineal puzzle is invalidated and we must ask different questions about men’s involvement in household economics (see also Macfarlan et al., this issue). This does not discount men’s involvement in all societies, but speaks to the need to quantify parental and allo-parental investments in different contexts.

## 4.0 The Expendable Male Hypothesis

> “…most students of the evolution of mating systems experienced a strong intuitive commitment to the idea that females were handicapped without male help.” (Gowaty 1996: 488)
>
> “…the males [of multi-male, multi-female primate societies] have apparently evolved to maximize matings, accepting a low confidence of paternity and showing less parental care than in other social groups” (Alexander 1974: 331)

Based on the foregoing discussion, the expendable male hypothesis posits as its fundamental premises that the matrilineal puzzle: (i) does not exist; (ii) that males do not decide whether to invest in nieces or nephews *or* in their own biological children; (iii) rather, males *either* invest little in parenting on the whole *or* they invest more intensely in their own biological children, when paternity is more certain; and (iv) that male investments in natal kin may be motivated less by genetic relatedness and more by (a) other benefits reaped through cooperation (e.g., reciprocity), or (b) constraints that promote cooperative breeding among natal kin (e.g., ‘helpers at the nest’/ecological constraints). Finally, (v) matriliny prevails in environments in which women’s efforts are sufficient to meet their families’ needs and in which men do not benefit disproportionately from usurping control of resources from women (e.g., because mating is a more viable strategy). Rather, in some environments, women’s investment in conjugal families may present unacceptable returns if ecologically expensive men [126,127] put household welfare at risk [128], and men may opt for strategies that do not involve significant effort being invested in any household.

Many aspects of matrilineal and matrifocal kinship may disincentivize paternal investment in biological offspring. Male parental investment is likely when a) paternity is relatively certain, b) opportunity costs of care are low, and c) paternal investment enhances men’s fitness more than investments in other components of fitness [112,129–132]. It has already been established that paternity certainty may be low for many males in matrilineal systems, whether due to protracted absences (e.g., warfare), residence systems such as natal philopatry that prevent effective mate guarding [cf. 133], etc. Another perspective on paternity is that, in the absence of strong investments into offspring, men may have little *interest* in whether paternity is assured. Ecologies [59,118] and institutions [12,63,67] that limit or decrease the impact of men’s investments on offspring fitness are as likely to lead to low paternity certainty as paternity certainty causing limited investment in offspring [86, see 134]. Mating markets are undoubtedly broader in such contexts, as women balance the benefits of multiple mating against fidelity [36,85] and men pursue mating opportunities that face little competition from parental responsibilities. We do not wish to imply that matriliny entirely precludes paternal investment – we have shown previously that in one economically transitional matrilineal context, fathers play important roles [129, see also 132]. Rather, we contend that support of childcare by males is often limited due to constraints that make such investments less beneficial than other behavioral strategies, for example mating or reciprocity-based contributions to the natal household. This broader perspective on human kinship configurations is consistent with patterns of parental investment across taxa: context-specific costs and benefits of caring for offspring generate differences in maternal and paternal investment both among species and among populations of the same species [15].

Consistent with this view are observations that when men have control over resources or make significant investments in childcare, such resources are used *not* to benefit their natal kin, but to advance their own reproductive agendas. Ethnographers have noted tendencies of matrilineal men to invest in their biological children, suggesting that men will act to maximize their own inclusive fitness even when doing so contradicts stated norms. In the Trobriand Islanders, for example, Malinowski noted a ‘strong tendency of surreptitious patrilineal transmission of property and influence’ [cited in 12]. Among the Nayar, who, like the Mosuo, were famous for a near-complete absence of fathers and husbands, Gough reported that ‘a man is said to have been especially fond of a child whom he knew with reasonable certainty to be his own’ [Gough 1961a:364, cited in 12]. Richards [3] observed that ‘in many matrilineal societies, a man typically must give his heritable possessions such as land to his sister’s children but is more likely to be involved in the day-to-day care of his putative offspring’ and that ‘men of wealth and distinction are able to reverse the usual rules of residence’. Pre-Communist Mosuo also illustrate this tendency: against a backdrop of middle- and lower-class matrilineal horticulturalists, upper-class families investing significant resources in their wives and children were patrilineal [120,135]. We are aware of limited quantitative evidence addressing men’s investments in nieces and nephews versus in their spousal households, but note that Wu and colleagues’ [34] results reveal that even unmarried Mosuo men invest almost as much labor in their spousal households as in their natal households and that married Mosuo men were more likely to be observed working on any farm than unmarried men. We suspect that this may be due to increasing demands on men’s labor by their affinal households [11,124, 134].

But why do men labor in their natal households at all? What do they do for their natal households? If women’s returns on investment in relationships with men are highly variable, and if the gendered division of labor requires any male input, then investment in male kin (brothers and uncles) could yield a higher inclusive fitness return than investment in a conjugal union. Men also provide an inclusive fitness benefit to the natal household that is unrelated to labor: mating. Ruth Mace’s research group has argued extensively that reproductive conflict underlies patterns of cooperation among the Mosuo, where competition among resources is stronger for women than men [34,77,124,136], and have pointed to the benefits of male mating within this context, speculating that “mothers may help adult sons stay in the natal household to enable them to mate more freely and/or do relatively less work than if they moved into their wife’s household.” [124]. In this view, matrilineal men assume roles that are potentially analogous to the drone in honey bee society, providing relatively little in the way of labor, but delivering significant benefits if successful on the mating market.

But surely, matrilineal men are less idle than honey bee drones? Male effort varies over the lifespan and lies on a spectrum from very little to very great [137]. Indeed, although we have often seen men being idle in our respective field sites, we have also observed men being quite helpful, including within their natal households [see also 34,129]. Even unmarried rural Caribbean men, who spend significantly less time working than do married men or even their pre-adolescent nieces, periodically take on extremely energy-intensive tasks requiring considerable upper body strength, such as forest clearing, which can be a substantial benefit to the natal household. Alexander [79] proposed a number of other under-appreciated factors motivating avuncular investments in nieces and nephews: the net benefit of investment in nieces and nephews increases “if the males involved are: (1) young and childless brothers, (2) brothers estranged from other wives, (3) unusually wealthy or powerful brothers, or (4) brothers who are for other reasons unlikely to be successful with their own offspring”. Such males experience relatively little conflict between investing in their own offspring and investing in their siblings’ offspring. Note that male effort in their natal households need not relate directly to allo-care of sisters’ children to be beneficial – men may, for example, be underwriting the costs of their own subsistence in the household [138,139] or providing labor in return for reciprocal benefits delivered from other members of the household. Thus, although we may need to seek explanations that venture beyond kin selection and paternity certainty to understand the variation in male investments [63,140], we may not require new explanations to solve a puzzle that does not exist.

As a final point of the expendable male hypothesis, we argue that the variability in male investments in their natal and affinal households in matrifocal societies is made possible by a labor force that largely consists of female kin [101], including maternal grandmothers, aunts, adolescent girls [49,115,127,141–143]. This labor force provides a sufficient basis for subsistence and supports the development of male and female descendants. Furthermore, the ability of women to provide the resources necessary to support their descendants disincentivizes additional male investment because it decreases offspring sensitivity to paternal investments in children [see ‘The Female Competence Hypothesis’ in 1]. This argument reignites the idea that the relative contributions of men versus women to subsistence plays a role in determining whether kinship systems are female-or male-biased [12,68,144–146] by considering how female contributions to subsistence impact male motivations to provide additional care [1,147]. Significant support from affinal men is helpful in certain contexts (e.g., when children become very expensive to rear)[148], but is not necessary to meet basic subsistence needs in matrifocal contexts [cf. 113]. Rather, as male support becomes necessary, kinship systems are anticipated to transition toward patrilineal/virilocal (if assistance from kin is required) or nuclear (if households are self-sufficient) [11,134].

| Hypothesis | Prediction | Empirical Support |
| --- | --- | --- |
| 1. Paternity certainty (PC) | 1. PC inversely associated with matriliney | Broad support for prediction, but PC alone is often considered inadequate |
| 2. MDBI/ Resources | 1. Expansive or labor-limited resource bases are associated with matriliney<br>2. Diminished PC and shallow RS/resource slope for men favor daughter-biased investment<br>3. Under matrilineal inheritance, uncles might transfer resources to sororal nephews<br>4. Under polyandry, men might invest in sororal nieces and nephews | Broad comparative support; quantitative support in Mosuo, Chewa, Gabbra |
| 3. Yield to labor | 1. Complex: Diminishing returns to labor more likely to generate avuncular investment than linear returns | No empirical tests |
| 4. Expendable males | 1. Men do not invest in nieces and nephews at the expense of own children<br>2. Men's investments in nieces and nephews are often trivial and more likely when such investments involve few opportunity costs<br>3. Male investments in sisters' households offset the costs of support of males in those households<br>4. Non-trivial investments are biased toward own children<br>5. (a) Pair-bonds are relatively fluid and (b) female mate-choice emphasizes indicators of genetic | Certain support from ethnographic anecdotes summarized in main text; Limited quantitative evidence; Starkweather and Hassan (this issue) support prediction 2<br>Extensive ethnographic support for prediction 5a. |
|  | quality over willingness and ability to provision.<br>6. Kinsmen's contributions to households are variable and substitutable by kinswomen or other exchanges such as hired labor.<br>7. Diminished PC and shallow RS/resource slope for men favor limited strategies that are decoupled from resources for men<br>8. Male control over resources is associated with investment in biological offspring |  |

## 5.0 Summary and Conclusion

We offer an initial articulation of the expendable male hypothesis and propose that this hypothesis offers an elaboration of existing solutions for the ‘matrilineal puzzle’ by denying that it is a puzzle and proposing instead that matriliny can evolve and remain stable in particular social and ecological circumstances. This significantly builds on prior scholarship, both evolutionary and non-evolutionary, in examining the relative costs and benefits of alternative strategies for men in different contexts. The expendable male hypothesis builds on existing scholarship by (i) grounding the expectation that males *either* invest little in their natal households to offset costs of belonging to a household, *or* they invest significantly in children to whom paternity is reasonably assured and whose wellbeing they can significantly improve (e.g., when male investments appreciably enhance female care). The hypothesis (ii) further predicts that males *do not invest significantly in nieces/nephews* at the expense of investing in their own children. Rather, we argue that established evolutionary theories of parental investment, cooperative breeding, and kin selection can explain variability in human male investments. The expendable male hypothesis therefore connects human matriliny to matriliny in other gregarious mammals, among whom male help is context-dependent, and almost never primarily focused on avuncular support of nieces and nephews. In these species, ecological constraints can render male investment in their own offspring either not possible[15] or not particularly beneficial to offspring success [149], so selection favors investment (typically more limited) in natal kin.

Our argument is compatible with many postulates of the matriliny-as-daughter-biased investment (MDBI) hypothesis [45,59], but our focus is different. Indeed, the same factors that bias investment towards daughters (i.e., low returns to investing resources in males) would also leave men in positions where their efforts in childrearing are largely substitutable by female kin [see also 150]. In such situations, men occupy tenuous positions as provisioners of their households, and often favor investment in other activities (e.g., mating). While the MDBI hypothesis leaves open the possibility that men make considerable investments in their children, but choose to bias those efforts toward daughters, we suggest instead that men might not invest significantly in many matrilineal contexts. In our view, MDBI could be thought of as a transgenerational consequence of expendable males. If men are relatively unimportant for childrearing and household wellbeing, then women are free to choose mates regardless of their mates’ willingness and ability to provision their households. Therefore, male fitness may be less sensitive to parental investment during development and more dependent on indicators of genetic quality. Hence, daughter-biased investment is evolutionarily rational when males are expendable.

The normative emphasis on men’s roles as mother’s brothers may seem to contradict our hypothesis. However, institutionalizing a special role for the mother’s brother does not tell us much about what such a role entails, and variability across societies belies a simple explanation [86]. Further, we emphasize that we do not discount all male investments in kin. Rather, we argue that such investments should arise when the opportunity costs of investments are minimal (e.g., because they are investing in limited or inconsistent ways, because they have no children of their own) or when such investments serve as a form of reciprocity between the mother and the mother’s brother. Institutionalizing the role of the mother’s brother may stabilize a relationship that, if inconsistent, is critical at specific junctures (e.g., protection from intruding men, acute investments of heavy labor). These efforts are compatible with dilute paternal investment in a man’s own offspring; they are only incompatible if they occur *at the expense of* a man’s own offspring. Significant paternity confusion might also be associated with the institutionalization of this role as men invest limited amounts across a number of known maternal descendants and putative heirs [36,151]. In any case, persistent emphasis on mother’s brothers, including in patrilineal societies [86,e.g., 152], suggests that men manage dual roles even in cases where they clearly favor their own biological children.

Thus, we argue, men’s behavior in matrilineal societies does not require special explanations beyond those applied to societies normatively organized around different principles of kinship. As discussed above, matrilateral investment in kin arises in numerous societies, even under strong patrilineal (even patriarchal) institutions or in WEIRD populations. The evidence for such investments is strongest for maternal grandmothers [4,e.g., 62,143,153], but has also been described for aunts and uncles [e.g., 75]. The particulars of such investment are often not described in relation to the strength of the investment (e.g., investing in college tuition versus providing treats) or to potential competing recipients, however. Our hypothesis highlights the critical need for more data that address investment decisions being made by men in a given context to define a) the level, nature, and timing of investment (e.g., trivial investments such as giving treats, b) the effects of given investments on children, and c) interests conflicting with investment in a particular child. Indeed, the life cycle of a male, regardless of society must also shift the costs and benefits of investing in different categories of kin as a man moves through phases of adolescence, marriage, and fatherhood [e.g., 66,154]. Recasting paternal investment decisions as the ‘matrilineal puzzle’ obscures the fact that similar decisions are simply being made in different contexts. Thus, the ‘matrilineal puzzle’ may be resolved if researchers investigated investment patterns in more detail in individual societies – rather than assuming ‘matriliny’ always comes with a certain set of behaviors.

We have intentionally disregarded the complexity of decisions that males and females make in relation to each other across different socio-ecologies [65] and have thereby presented what is undoubtedly an overly simplistic characterization of men as ‘expendable’ where there efforts are limited, inconsistent, unnecessary, substitutable, or applied to mating rather than parenting. We do not intend to trivialize men’s positions in webs of matrilineal kinship, but to question the assumptions that men are always in positions of authority that somehow require significant investments in sororal nieces and nephews at the expense of a man’s own children. We welcome theoretical refinement and robust empirical tests of the expendable male hypothesis.

## 6.0 Acknowledgements

The authors thank Monique Borgerhoff Mulder, two anonymous reviewers, and Laura Fortunato for helpful comments on the manuscript. Lingering problems are due to insufficient parenting by our fathers and mothers’ brothers. SM thanks the University of New Mexico ADVANCE team for supporting a workshop in which these ideas were presented and refined. Fieldwork among the Mosuo has been supported by the National Science Foundation (BCS 0717918 and 1461514), the University of Washington Chester Fritz and Foreign Language Area Studies Fellowships, and the American Philosophical Society. SM thanks Mosuo participants for tolerating her nosiness despite a lack of immediate returns to fitness. She also thanks her husband for not being expendable and Donna Leonetti for challenging her to think critically about the centrality of females to the evolution of human sociality. Adam Reynolds assisted in proofing this manuscript and redrawing figures.

## Footnotes

1 Indeed, patriliny is probably more puzzling from a broad comparative perspective, particularly given apparently limited ability to recognize paternal kin [e.g., 38,c.f., 39].

2 Mother’s-brother-sister’s-son descent and inheritance are often taken to represent ‘matriliny’ in ethnographic and evolutionary literature. As clarified within the main text, the ‘problem’ is relegated to avuncular investment in nieces and nephews, not to matriliny. For example, matrilineal systems that transmit resources directly from parents to daughters are relatively easily explained [e.g., 3,45].

3 In evolutionary scholarship, mother’s-brother-sister’s-son wealth transmission is often taken to represent ‘matriliny’. Other forms of matrilineal inheritance (e.g., daughter-biased investment) are not problematic for evolutionary scholars, though they remain puzzling to many socio-cultural anthropologists because they still divide male authority between the natal and affinal (i.e., marital) lineages.

